# c-Fos enhances influenza virus replication by stabilizing the M2 protein and promoting autophagosome accumulation

**DOI:** 10.64898/2026.03.05.709812

**Authors:** Xiaonan Chen, Junyang Yan, Mengxue Li, Hanbin Liu, Daiqiang Lu, Yang Wang, Qiao Zhang, Feng Gao, Jiaojiao Peng

**Affiliations:** Institute of Molecular and Medical Virology, School of Medicine, Jinan University, Guangzhou, China; Key Laboratory of Viral Pathogenesis & Infection Prevention and Control, School of Medicine, Jinan University, Guangzhou, China; Guangdong Zocode Biotechnology Services Co., Ltd, Guangzhou, China; State Key Laboratory of Respiratory Disease, National Clinical Research Center for Respiratory Disease, National Center for Respiratory Medicine, Guangzhou Institute of Respiratory Health, the First Affiliated Hospital of Guangzhou Medical University, Guangzhou, China

**Keywords:** c-Fos, influenza virus, M2 protein, autophagy, viral replication

## Abstract

The influenza A virus (IAV) employs multiple strategies to hijack host cellular machinery for efficient replication. While autophagosome accumulation is known to promote IAV replication, and the viral matrix protein 2 (M2) ion channel protein plays essential roles in viral uncoating, assembly, and autophagy induction, the mechanisms regulating M2 stability remain incompletely understood. Furthermore, although c-Fos participates in the replication of various viruses, its function in IAV infection has not been characterized. Here, we identify c-Fos as a critical proviral host factor that enhances IAV replication through stabilizing M2 protein and promoting autophagosome accumulation. We demonstrate that IAV infection triggers M2-mediated cytosolic calcium elevation, which upregulates c-Fos expression. The induced c-Fos physically interacts with M2 and prevents its proteasome and lysosomal degradation, thereby increasing M2 accumulation. This stabilized M2 protein as a viral protein directly facilitates viral replication while simultaneously promoting autophagosome accumulation to further enhance the process. Our findings reveal a novel positive feedback loop in IAV infection, establishing c-Fos as a key regulator of IAV replication through M2 stabilization, and highlighting these interactions as potential therapeutic targets for antiviral intervention.

**IMPORTANCE:** Influenza A virus (IAV) remains a significant global health threat, causing substantial morbidity and mortality. Understanding virus-host interactions is crucial for developing antiviral strategies. Here, we uncover a previously unrecognized positive feedback loop in which IAV exploits the host transcription factor c-Fos to enhance its own replication. We demonstrate that IAV infection upregulates c-Fos through M2-mediated calcium signaling, and the induced c-Fos in turn stabilizes the viral M2 protein. This stabilized M2 not only directly supports viral replication but also promotes autophagosome accumulation, further facilitating virus production. Our findings establish c-Fos as a critical proviral host factor and reveal a mechanism by which IAV hijacks host cellular machinery to create a favorable environment for efficient replication. The identification of this c-Fos-M2 axis not only advances our understanding of IAV-host interactions but also opens new avenues for therapeutic intervention targeting this vulnerability in the viral life cycle.

## INTRODUCTION

Influenza A virus (IAV), a member of the Orthomyxovirus family, possesses a lipid envelope and a segmented, negative-sense, single-stranded RNA genome [1]. IAV remains a significant global health threat, causing seasonal epidemics and occasional pandemics due to its high mutation rate and zoonotic potential [2, 3]. The World Health Organization estimates that annual epidemics result in about 1 billion infections, 3-5 million cases of severe illness, and 290,000-650,000 respiratory deaths globally, underscoring its persistent public health burden [4]. Successful viral replication relies on the exploitation of host cellular machinery, with numerous viral and host factors participating in this intricate process. Among these, the IAV M2, an ion channel protein, plays essential roles in viral uncoating, assembly, and budding [5–7]. Additionally, M2 has been implicated in inducing autophagy, a conserved cellular process that viruses often hijack to support their replication [8, 9]. Despite its critical functions, the mechanisms regulating M2 protein stability remain incompletely understood, particularly how host factors modulate its turnover to facilitate viral propagation.

c-Fos is an important immediate early gene (IEG)-encoded transcription factor belonging to the AP-1 family, playing critical roles in diverse biological processes. It regulates cell proliferation and differentiation, and is rapidly induced in response to external stimuli such as oxidative stress, heat shock, and ultraviolet radiation, thereby mediating stress responses [10, 11] and facilitating cellular adaptation to environmental signals [12]. Within the immune system, c-Fos participates in the regulation of immune responses by modulating immune cell functions [13]. Interestingly, growing evidence suggests that c-Fos participate in host-virus interactions [14–17], though its specific role in IAV infection remains unexplored.

Sarco/endoplasmic reticulum Ca²⁺-ATPase (SERCA) pumps Ca²⁺ from the cytosol into the ER lumen, thereby maintaining low resting cytosolic calcium levels, inhibition of SERCA activity disrupts this homeostasis, which is known to induce ER stress and influence both viral infection and autophagy pathways [18–22]. Our previous research demonstrated that SERCA regulates influenza virus-induced autophagosome accumulation and viral protein expression [23]. In this study, through screening for key host factors mediating this regulation, we identified c-Fos as a critical mediator of SERCA’s effect on influenza virus and accordingly investigated its role and underlying mechanism in viral replication.

Here, we identify a positive feedback loop centered on c-Fos that potentiates influenza A virus replication. We demonstrate that M2-mediated calcium influx upregulates c-Fos, which in turn stabilizes the M2 protein by protecting it from proteasomal and lysosomal degradation. This stabilized M2 serves a dual function, as it directly facilitates viral replication and concurrently promotes autophagosome accumulation to further support viral propagation. Our findings reveal c-Fos as a critical proviral host factor exploited by IAV to sustain M2 levels and enhance autophagy, illuminating a previously unrecognized mechanism in the viral life cycle and identifying potential therapeutic targets for antiviral intervention.

## RESULTS

### c-Fos is a key host factor in SERCA-mediated regulation of IAV replication

We have previously shown that IAV infection inhibits sarcoplasmic/endoplasmic reticulum calcium ATPase (SERCA) activity and that SERCA knockdown enhances viral protein expression [23]. To further delineate the functional role of SERCA in IAV replication, we infected H1395 cells with two distinct influenza virus strains (PR8 and WSN) and assessed viral protein expression following SERCA knockdown or pharmacological activation with CDN1163. Transfection of SERCA-targeting small interfering RNA (siRNA) significantly increased the expression levels of IAV nucleoprotein (NP) (Fig. 1A), whereas SERCA activation with CDN1163 suppressed NP expression (Fig. 1B). These results collectively indicate that SERCA negatively regulates influenza virus replication.

**Fig 1.**
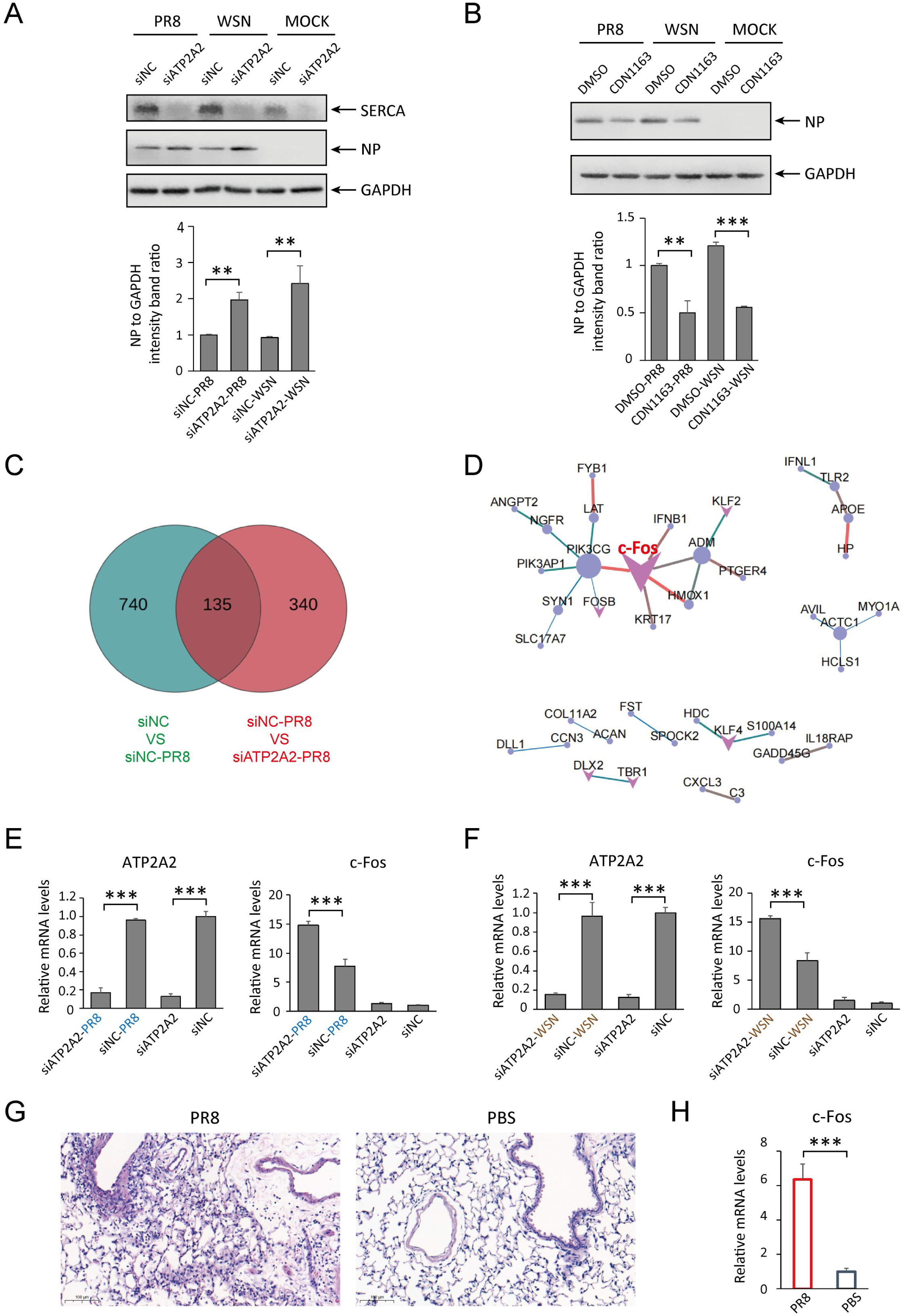
c-Fos is a key host factor in SERCA-mediated regulation of IAV replication. (A) H1395 cells were transfected with siNC or siATP2A2 for 24 h and then infected with PR8/WSN or mock infected. The cell lysates were subjected to immunoblotting analysis with the indicated antibody. (B) H1395 cells were infected with PR8/WSN or mock infected and then treated with CDN1163 or DMSO. The cell lysates were subjected to immunoblotting analysis with the indicated antibody. (C) The Venn diagram shows that green represents the number of genes significantly upregulated by siNC-PR8 relative to the siNC group, red represents the number of genes significantly upregulated by siATP2A2-PR8 relative to siNC-PR8, and the overlapping region in the middle represents the number of genes co-upregulated. (D) PPI network showing interactions among common target genes in Venn diagram. (E and F) H1395 cells were transfected with siNC or siATP2A2 for 24 h and then infected with PR8 (E) or WSN (F), and the transcription levels of ATP2A2 and c-Fos were monitored by qRT-PCR. (G) H&E staining of mouse lung tissue. Scale bar, 100 μm. (H) mRNA levels of c-Fos in the lung tissue of infected mice were determined by RT-qPCR. All data are presented as means ± SD from at least three independent experiments. Significance was determined by Student’s t test. **, P < 0.01; ***, P < 0.001.

To identify host factors mediating this regulatory effect, we performed transcriptomic analysis on SERCA-knockdown (siATP2A2) and control (siNC) cells, either mock-infected or infected with the PR8 IAV strain. Genes upregulated by PR8 infection (vs. mock infection) were intersected with those upregulated in siATP2A2-transfected, PR8-infected cells (vs. siNC-transfected, PR8-infected cells), yielding 135 candidate host factors (Fig. 1C). Protein-protein interaction (PPI) network analysis and pathway enrichment analysis of these candidate genes identified c-Fos as the most central host factor through which SERCA exerts its regulatory effect on IAV replication (Fig. 1D). We next validated the transcriptomic findings by quantitative real-time PCR (qPCR). Consistent with the RNA-seq data, IAV infection upregulated c-Fos transcription, and this inductive effect was further enhanced by SERCA knockdown (Fig. 1E). This observation was recapitulated using another influenza virus strain (WSN), which showed a similar trend: SERCA knockdown potentiated virus-induced c-Fos upregulation (Fig. 1F).

To further examine the effect of influenza virus on c-Fos expression in vivo, we intranasally inoculated mice with the PR8 strain and assessed c-Fos expression in lung tissues. Three days post-infection, mice were euthanized, and lung tissues were collected for analysis. Hematoxylin-eosin (HE) staining confirmed successful IAV infection and associated inflammatory responses in the lungs (Fig. 1G). Consistent with the in vitro results, IAV infection significantly upregulated c-Fos expression in vivo (Fig. 1H).

Taken together, these results demonstrate that c-Fos is a key host factor mediating SERCA-dependent regulation of influenza virus replication.

### c-Fos promotes IAV replication and virus-induced autophagosome accumulation

Given the evidence that IAV strongly induces c-Fos both in vivo and in vitro, and that SERCA negatively regulates both c-Fos and IAV replication, we hypothesized that c-Fos might play a functional role in influenza virus replication. To test this, we examined lung cell lines (H1395) and observed that infection with either PR8 or WSN influenza virus strains significantly upregulated c-Fos expression in H1395 cells, and RNA interference-mediated knockdown of c-Fos markedly suppressed viral gene expression in infected cells (Fig. 2A-2C). Furthermore, plaque assays revealed that c-Fos knockdown reduced viral titers by 10-fold (Fig. 2D) and 14-fold (Fig. 2E) compared to control cells. Consistent results were obtained in another lung cell line, H1975 cells (Fig. 2F-2I). Together, these findings demonstrate that c-Fos is a critical host factor for efficient influenza virus replication.

**Fig 2.**
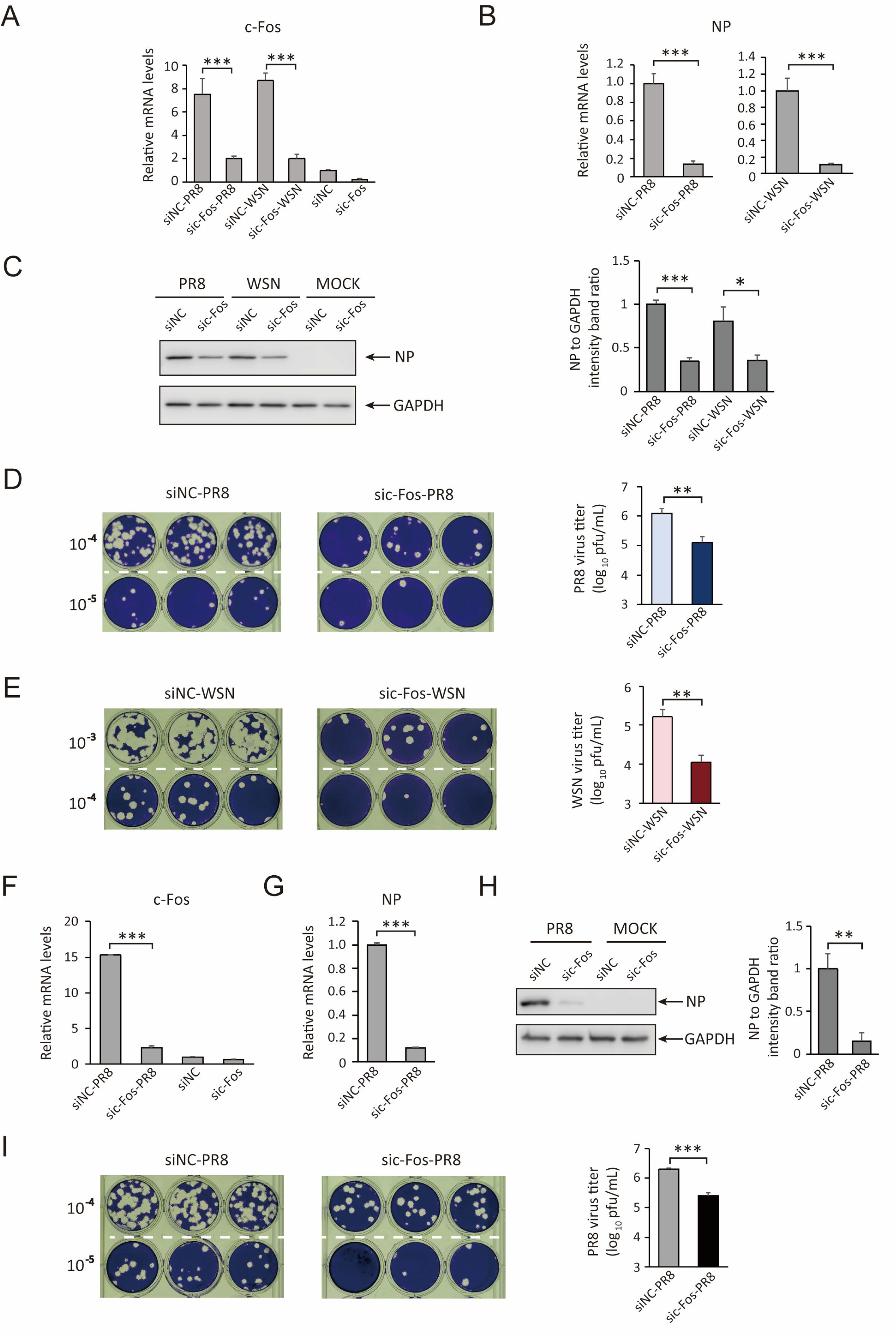
c-Fos promotes IAV replication. (A–E) H1395 cells were transfected with siNC or sic-Fos for 24 h, then either mock-infected or infected with PR8 or WSN virus. (A, B) qRT-PCR analysis of c-Fos (A) and viral NP (B) mRNA levels. (C) Immunoblot analysis of cell lysates using the indicated antibodies. (D, E) Viral titers in supernatants collected 24 h post-infection with PR8 (D) or WSN (E), determined by plaque assay. Serial dilutions of viral supernatants are shown on the left, with three replicates per dilution. (F–I) H1975 cells were transfected with siNC or sic-Fos for 24 h, then mock-infected or infected with PR8 virus. (F, G) qRT-PCR analysis of c-Fos (F) and viral NP (G) mRNA levels. (H) Immunoblot analysis of cell lysates using the indicated antibodies. (I) Viral titers in supernatants collected 24 h post-infection, determined by plaque assay. Serial dilutions are shown on the left, with three replicates per dilution. All data are presented as means ± SD from at least three independent experiments. Significance was determined by Student’s t test. *, P < 0.05; **, P < 0.01; ***, P < 0.001.

Since our previous studies demonstrated that SERCA regulates influenza virus-induced autophagy [23], we investigated whether c-Fos, a downstream target of SERCA, modulates this process. In PR8-infected H1395 and H1975 cells, immunofluorescence staining for the viral NP protein and the autophagosome marker LC3B showed that influenza infection robustly induced autophagosome accumulation (Fig. 3A), consistent with prior findings. Notably, c-Fos knockdown in both cell lines significantly suppressed this virus-induced autophagosome production (Fig. 3A). To quantify this effect, we measured the conversion of LC3-I to LC3-II, a key step in autophagosome formation. Immunoblot analysis revealed that the LC3-II/ACTB ratio was significantly increased in PR8-infected cells compared to uninfected controls. Conversely, this ratio was significantly reduced upon c-Fos knockdown in PR8-infected cells (Fig. 3B), confirming the positive regulatory role of c-Fos in influenza-triggered autophagy. These observations were recapitulated using the WSN viral strain, demonstrating consistent effects across different influenza variants. Collectively, our results establish c-Fos as a promoter of influenza virus-induced autophagosome accumulation.

**Fig 3.**
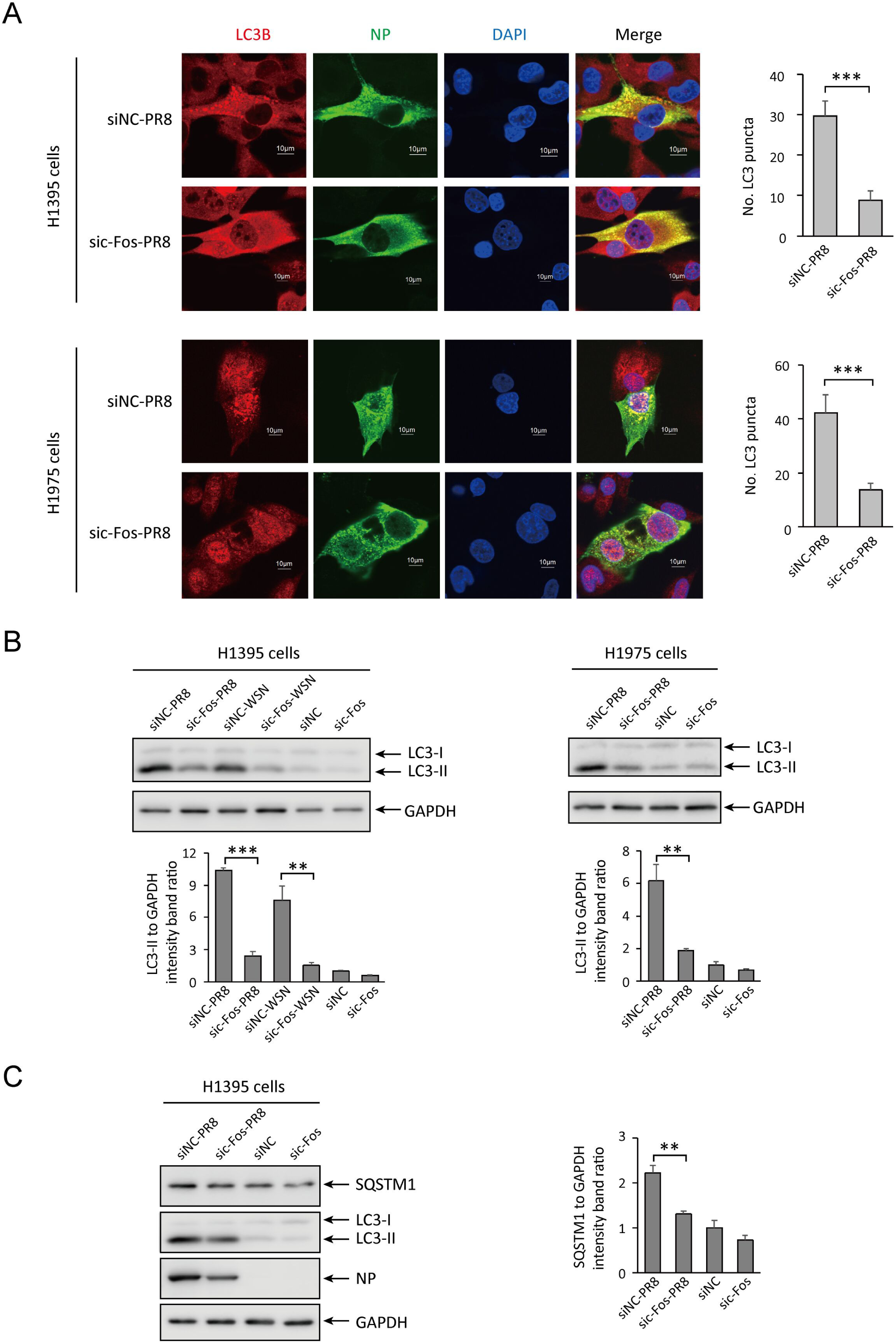
c-Fos enhances IAV-induced autophagosome accumulation. (A) Following a 24 h c-Fos knockdown and subsequent PR8 virus infection in H1395 and H1975 cells, then cells were fixed, immunostained with antibodies against LC3 and NP, and examined using confocal microscopy. Scale bar, 10 μm. Quantification of the number of LC3 puncta per cell were measured using a minimum of 10 images per condition. (B) H1395 or H1975 cells were transfected with siNC or sic-Fos for 24 h and then infected with PR8, WSN or mock infected. The cell lysates were subjected to immunoblotting analysis with the indicated antibody. (C) H1395 cells were transfected with siNC or sic-Fos for 24 h and then infected with PR8 or mock infected. The cell lysates were subjected to immunoblotting analysis with the indicated antibody. All data are presented as means ± SD from at least three independent experiments. Significance was determined by Student’s t test. **, P < 0.01; ***, P < 0.001.

To determine whether c-Fos promotes influenza virus-induced autophagosome accumulation by enhancing their generation or inhibiting their degradation, we assessed the level of SQSTM1. Since SQSTM1 is degraded when autophagic flux is complete, its protein level serves as an indicator of autophagic flux efficiency. Our results showed that infection with influenza virus PR8 upregulates SQSTM1 (Fig. 3C), and knockdown of c-Fos downregulated SQSTM1 (Fig. 3C), suggesting that c-Fos depletion alleviates the blockade of autophagic flux caused by influenza virus. Based on these findings, we conclude that knockdown of c-Fos inhibits influenza virus-induced autophagosome accumulation by promoting the autophagic flux process.

### IAV promotes c-Fos expression by inducing cytosolic calcium elevation via its M2 protein

Our previous studies showed that influenza virus infection inhibits SERCA activity [23], a calcium pump critical for pumping calcium ions from the cytosol into the endoplasmic reticulum [22], and that SERCA knockdown enhances IAV-induced c-Fos upregulation (Fig. 1E, 1F). Based on these findings, we hypothesized that IAV elevates c-Fos expression by increasing cytosolic calcium levels. To test this, we first measured cytosolic calcium changes upon infection with PR8 or WSN strains using the fluorescent indicator Fluo-4. Flow cytometry analysis revealed increased Fluo-4 fluorescence in both PR8- and WSN-infected cells (Fig. 4A), indicating that IAV infection elevates cytosolic calcium. To determine whether this calcium flux is responsible for c-Fos induction, we treated cells with BAPTA-AM (12 µM, a non-cytotoxic concentration; Fig. 4B) to chelate cytosolic calcium. Blocking calcium elevation markedly reduced IAV-triggered c-Fos expression (Fig. 4C), supporting our hypothesis that IAV upregulates c-Fos through calcium signaling.

**Fig 4.**
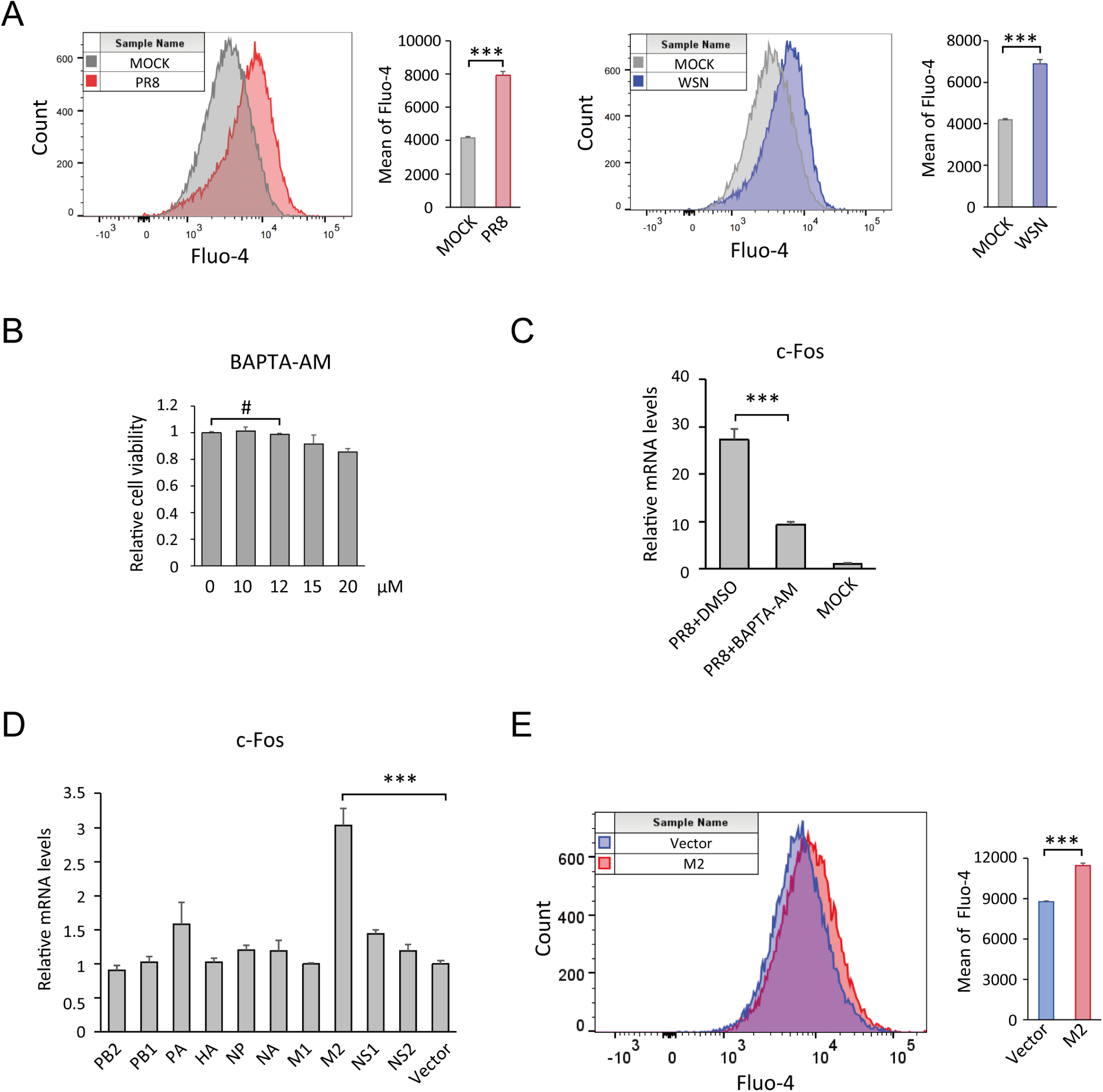
IAV promotes c-Fos expression by inducing cytosolic calcium elevation via its M2 protein. (A) H1395 cells were infected with PR8, WSN or mock infected. Cytosolic Ca²⁺ was labeled with Fluo-4, and fluorescence intensity was measured by flow cytometry, with 30,000 events collected per sample for analysis. (B) Cell viability was assessed by CCK-8 assay in BAPTA-AM treated cells. (C) H1395 cells were pretreated with BAPTA-AM or Mock for 2 h, followed by PR8 virus infection. At 24 h post-infection, c-Fos transcription levels were measured by qRT-PCR. (D) 293H cells were transfected with plasmids encoding individual viral proteins of IAV PR8 strain (empty vector as control). Cells were harvested 48 h post-transfection, and c-Fos transcription levels were analyzed by qRT-PCR. (E) 293H cells were transfected with PR8-M2 plasmid or empty vector. After 48 h, cytosolic Ca²⁺ was labeled with Fluo-4, and fluorescence intensity was measured by flow cytometry, with 30,000 events analyzed per sample. All data are presented as means ± SD from at least three independent experiments. Significance was determined by Student’s t test: ***, P < 0.001; #, P > 0.05.

We next sought to identify the viral factor mediating this effect. By individually expressing each of the 10 IAV proteins and assessing their ability to induce c-Fos, we found that the M2 protein alone was sufficient to drive c-Fos transcription (Fig. 4D). M2 expression also elevated cytosolic calcium levels (Fig. 4E), suggesting that M2 promotes c-Fos expression specifically through calcium signaling.

Having established that c-Fos suppression inhibits IAV replication and autophagy (Fig. 3), we next sought to determine whether BAPTA-AM could phenocopy this effect by blocking calcium-dependent c-Fos upregulation. Indeed, chelating cytosolic calcium not only reduced c-Fos levels but also attenuated viral replication, as measured by viral protein and titer levels (Fig. 5A-5C), and suppressed autophagosome accumulation, as assessed by LC3 marker analysis (Fig. 5C).

**Fig 5.**
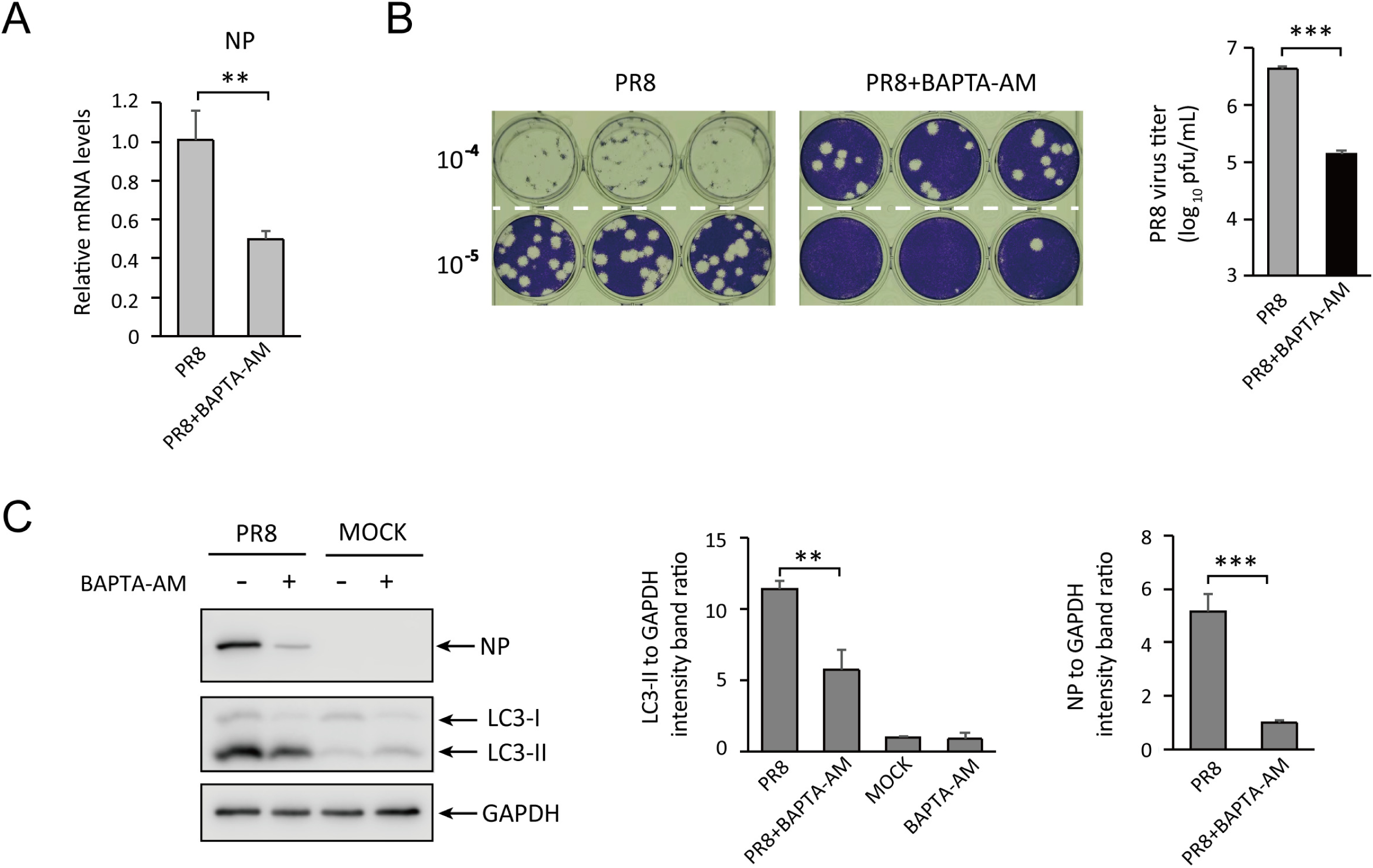
Chelation of cytosolic calcium suppresses IAV replication and virus-triggered autophagy. (A) H1395 cells were pretreated with BAPTA-AM or DMSO for 2 h, followed by PR8 virus infection. NP mRNA levels were quantified by qRT-PCR at 24 h post-infection. (B) Viral titers in the culture supernatants were determined by plaque assay at 24 h post-infection. The left panel indicates the dilution factor of the supernatant, with three replicates per dilution. (C) H1395 cells were pretreated with BAPTA-AM or Mock for 2 h, followed by PR8 virus or mock infection, cell lysates were analyzed by immunoblotting with the indicated antibodies. All data are presented as means ± SD from at least three independent experiments. Significance was determined by Student’s t test: **, P < 0.01; ***, P < 0.001.

Collectively, these results demonstrate that IAV promotes c-Fos expression by inducing cytosolic calcium elevation through its M2 protein.

### c-Fos stabilizes IAV M2 protein by blocking proteasomal and lysosomal degradation pathways

As c-Fos is a known transcription factor, we first investigated whether its enhancement of IAV replication is dependent on its transcriptional activity. Using T5224, a specific inhibitor of this activity, we pretreated cells with non-cytotoxic concentrations (10 µM and 20 µM) and observed no significant effect on IAV NP protein expression (Fig. 6A). This result indicates that c-Fos promotes IAV replication through a transcription-independent mechanism. Since we previously identified that IAV upregulates c-Fos expression via its M2 protein, we next examined whether c-Fos reciprocally regulates M2 protein. Co-transfection experiments with Flag-tagged c-Fos and PR8-M2 plasmids revealed that c-Fos significantly increased M2 protein levels without affecting M2 mRNA expression (Fig. 6B and 6C). This observation led us to hypothesize that c-Fos might interact with M2 to prevent its degradation. Co-immunoprecipitation (Co-IP) assays confirmed a specific interaction between c-Fos and M2 protein (Fig. 6C), while other viral proteins such as NP showed neither binding to c-Fos nor any effect on its abundance (Fig. 6D). Supporting these findings, confocal microscopy revealed clear co-localization of c-Fos with M2 in cells (Fig. 6E). Additionally, cycloheximide (CHX) chase analysis revealed that c-Fos overexpression prolonged the half-life of M2 (Fig. 6F). These results collectively indicate that c-Fos stabilizes M2 protein by inhibiting its degradation.

**Fig 6.**
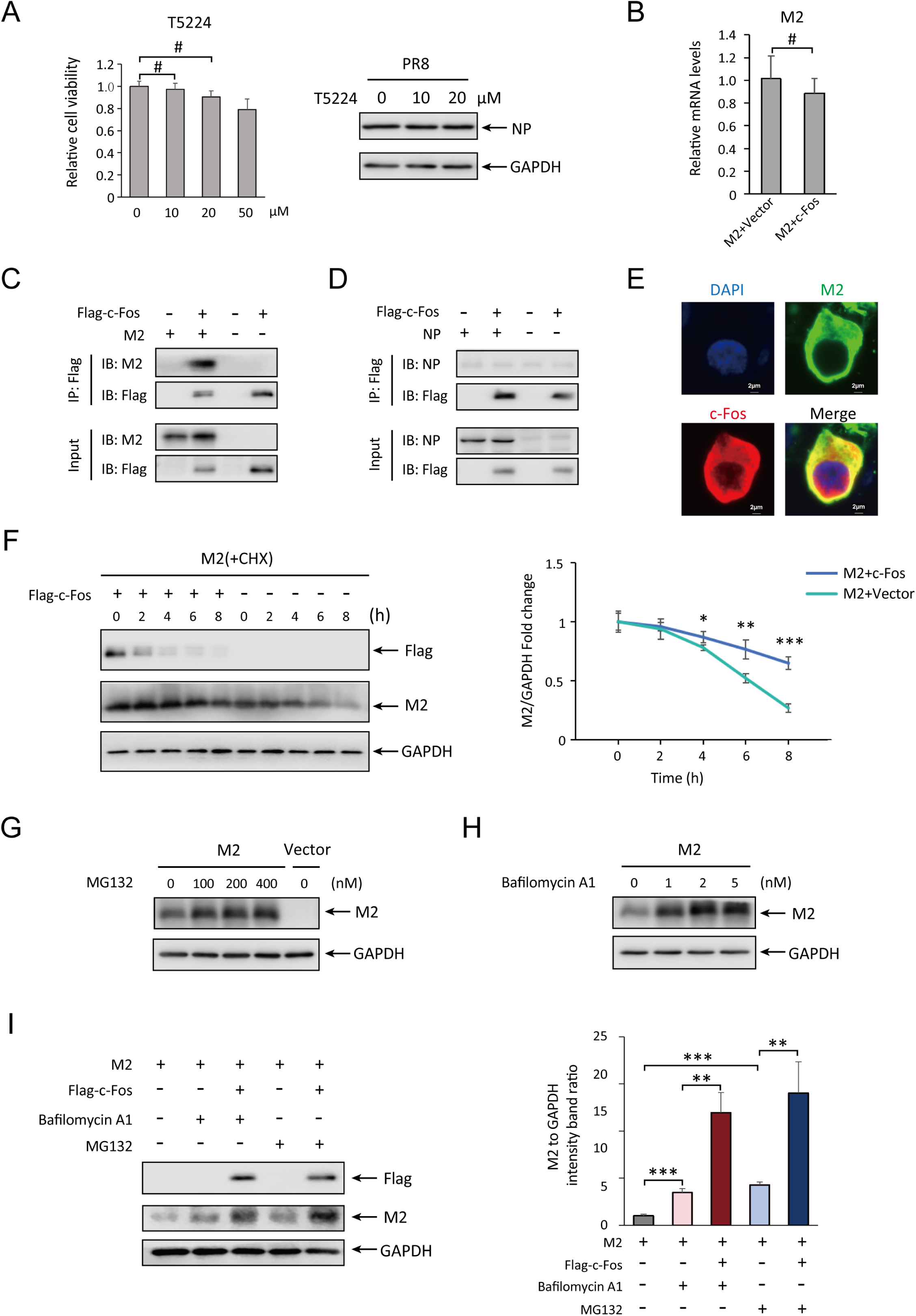
c-Fos stabilizes IAV M2 protein by blocking proteasomal and lysosomal degradation pathways. (A) Cell viability was assessed by CCK-8 assay following T5224 treatment. For infection experiments, cells were pretreated with 0, 10, or 20 μM T5224 for 2 h before PR8 virus infection. Protein lysates were harvested at 24 h post-infection and analyzed by Western blot using anti-NP and anti-GAPDH antibodies. (B) 293H cells were co-transfection with M2 and c-Fos plasmids, alongside M2 and vector controls. After 48 h, cells were lysed and RNA was extracted for RT-qPCR analysis of M2 mRNA expression. (C and D) Co-immunoprecipitation and immunoblot analysis of 293H cells co-transfected with Flag-c-Fos and M2 (C) or NP (D). (E) 293H cells were co-transfection with Flag-c-Fos and M2 for 48 h, followed by immunostaining for Flag (red) and M2 (green). Representative confocal immunofluorescence images are shown. Scale bars, 2 μm. (F) 293H cells were co-transfected with M2 and Flag-c-Fos or empty vector. Cells were treated with 50 μg/mL CHX and harvested at the indicated time points (0 - 8 h). M2 protein levels were analyzed by Western blot. (G and H) 293H cells were transfected with M2 or empty vector. At 6 h post-transfection, cells were treated with MG132 (100, 200, 400 nM), Bafilomycin A1 (1, 2, 5 nM), or DMSO control. After 48 h, M2 protein expression was assessed by Western blot. (I) 293H cells were co-transfected with M2 and Flag-c-Fos or vector, followed by treatment with MG132 (200 nM) or Bafilomycin A1 (2 nM) for 48 h. M2 protein levels were analyzed by Western blot. All data are presented as means ± SD from at least three independent experiments. Significance was determined by Student’s t test: *, P < 0.05; **, P < 0.01; ***, P < 0.001. #, P > 0.05.

To determine how c-Fos regulates the degradation of M2 protein, we first investigated the pathways responsible for M2 degradation. Using the proteasome inhibitor MG132 and the lysosome inhibitor Bafilomycin A1, we confirmed that M2 is degraded through both the proteasomal and lysosomal pathways (Fig. 6G and 6H). Next, to elucidate whether c-Fos inhibits M2 degradation via these pathways, we co-transfected c-Fos and M2 while separately blocking each degradation mechanism. When proteasomal degradation was inhibited by MG132, c-Fos overexpression further suppressed M2 degradation, suggesting that c-Fos blocks lysosomal degradation of M2 (Fig. 6I). Similarly, our results revealed that c-Fos also inhibits the proteasomal degradation of M2 (Fig. 6I).

Taken together, these results demonstrate that c-Fos enhances M2 protein stability by simultaneously inhibiting both its lysosomal and proteasomal degradation pathways.

### c-Fos-M2 axis enhances autophagy to promote viral replication

Building upon our previous demonstration that c-Fos promotes both IAV replication and virus-induced autophagosome accumulation, we sought to elucidate the underlying mechanism. Since M2 is not only essential for IAV replication but has also been shown to be the key viral protein driving autophagosome accumulation, and given the established role of autophagy in promoting influenza infection, we hypothesized that c-Fos enhances IAV replication by stabilizing M2 protein, thereby promoting autophagosome accumulation. To test this hypothesis, we first confirmed M2’s specific role in IAV-induced autophagy in H1395 cells. Expression of all 10 IAV proteins individually revealed that only PR8 M2 protein could independently increase levels of the autophagosome marker LC3-II (Fig. 7A). Importantly, c-Fos overexpression enhanced both M2 protein levels and LC3-II accumulation, while showing no such effect on IAV NP protein (Fig. 7B and 7C). Conversely, RNAi-mediated knockdown of c-Fos reduced both M2 protein stability and LC3-II levels (Fig. 7D). These results demonstrate that c-Fos promotes autophagosome accumulation specifically through M2 stabilization.

**Fig 7.**
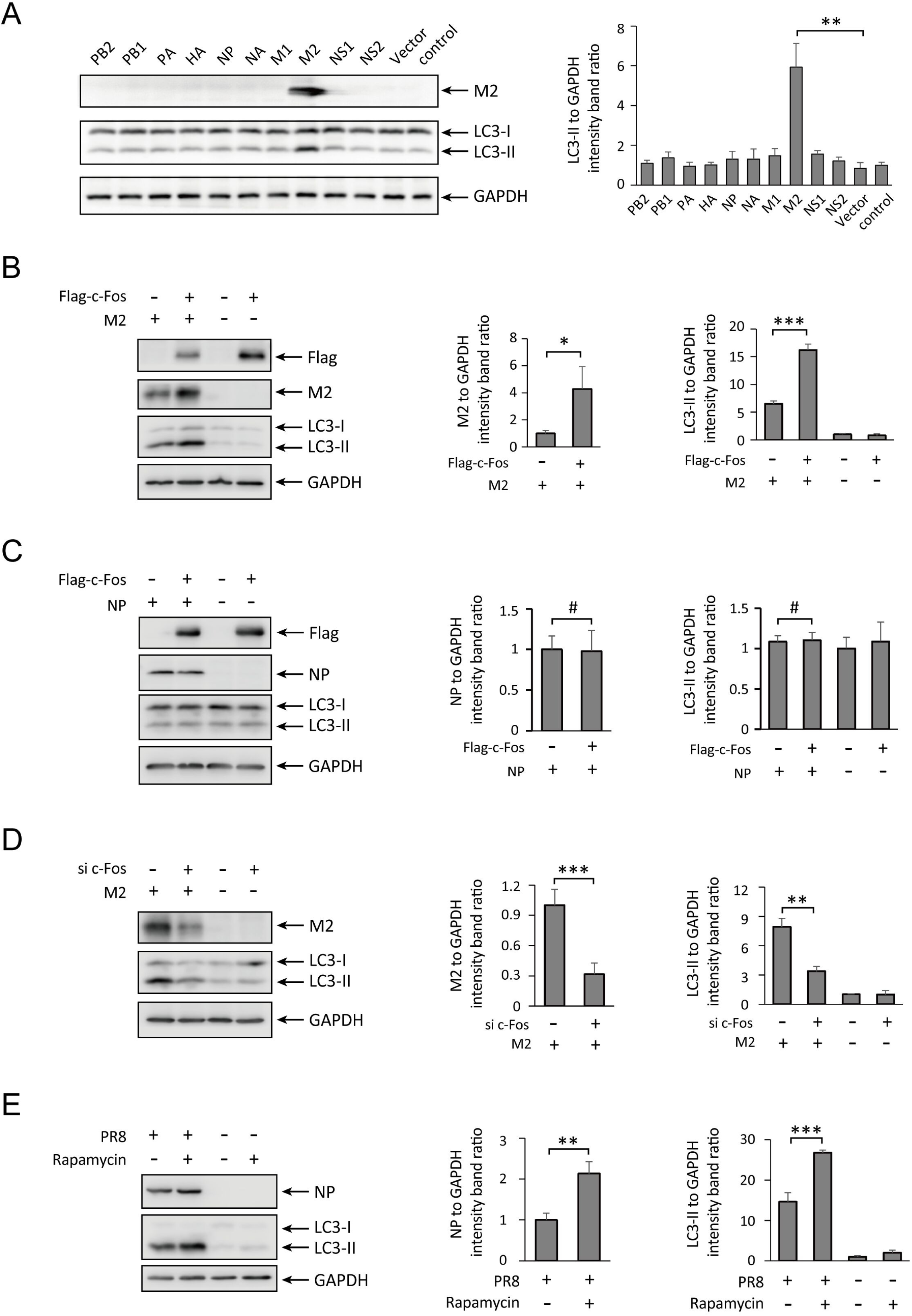
c-Fos promotes IAV replication by stabilizing M2 protein to induce pro-viral autophagy. (A) 293H cells were transfected with plasmids encoding individual viral proteins of the IAV PR8 strain, empty vector was used as a negative control. Cells were harvested 48 h post-transfection, and lysates were analyzed by Western blotting to assess LC3 lipidation (LC3-II) levels, with GAPDH used as a loading control. (B and C). 293H cells were co-transfected with Flag-c-Fos and M2 (B) or NP (C) plasmids, empty vector was used as a control. Cells were harvested 48 h post-transfection and analyzed by Western blotting for the indicated proteins. (D) 293H cells were transfected with siRNA targeting c-Fos (sic-Fos) or negative control siRNA (siNC). At 24 h post-siRNA transfection, cells were further transfected with M2 plasmid and harvested 48 h later. Protein levels were analyzed by Western blotting. (E) H1395 cells were pretreated with 100 nM Rapamycin or an equivalent volume of DMSO for 6 h, followed by infection with PR8 virus (MOI=1) or mock infection. Cells were harvested 12 h post-infection and analyzed by Western blotting for viral NP and LC3-II levels. All data are presented as means ± SD from at least three independent experiments. Significance was determined by Student’s t test: *, P < 0.05; **, P < 0.01; ***, P < 0.001. #, P > 0.05.

To establish the functional link between M2-induced autophagy and viral replication, we treated cells with the autophagy inducer rapamycin prior to IAV infection. Enhanced autophagosome accumulation significantly increased viral replication (Fig. 7E). Taken together, our findings reveal that c-Fos stabilizes the M2 protein by inhibiting its degradation, which subsequently enhances virus-induced autophagosome formation, thereby promoting influenza virus replication.

### Interaction sites of c-Fos-M2 and their potential as therapeutic targets for anti-IAV drug development

To explore the potential of targeting the c-Fos-M2 interaction for antiviral therapy, we predicted their binding sites and assessed the sequence conservation of these sites within the viral M2 protein. Molecular docking analysis identified several potential interaction sites on M2 for c-Fos, including ARG-18, ASN-20, SER-23, ASP-24, ASP-44, LYS-49, TYR-57, GLU-66, SER-71, and GLU-75 (Fig. 8A). We then retrieved 16,532 complete H1N1, 2,813 H1N2, 215 H2N2, and 16,918 H3N2 human IAV M2 protein sequences from the UniProt database. These sequences were grouped by subtype, and the conservation of each amino acid was analyzed. Among these, SER-23, ASP-24, ASP-44, LYS-49, GLU-66, SER-71, and GLU-75 exhibit high conservation across different subtypes (indicated by arrows in the analysis) (Fig. 8B).

**Fig 8.**
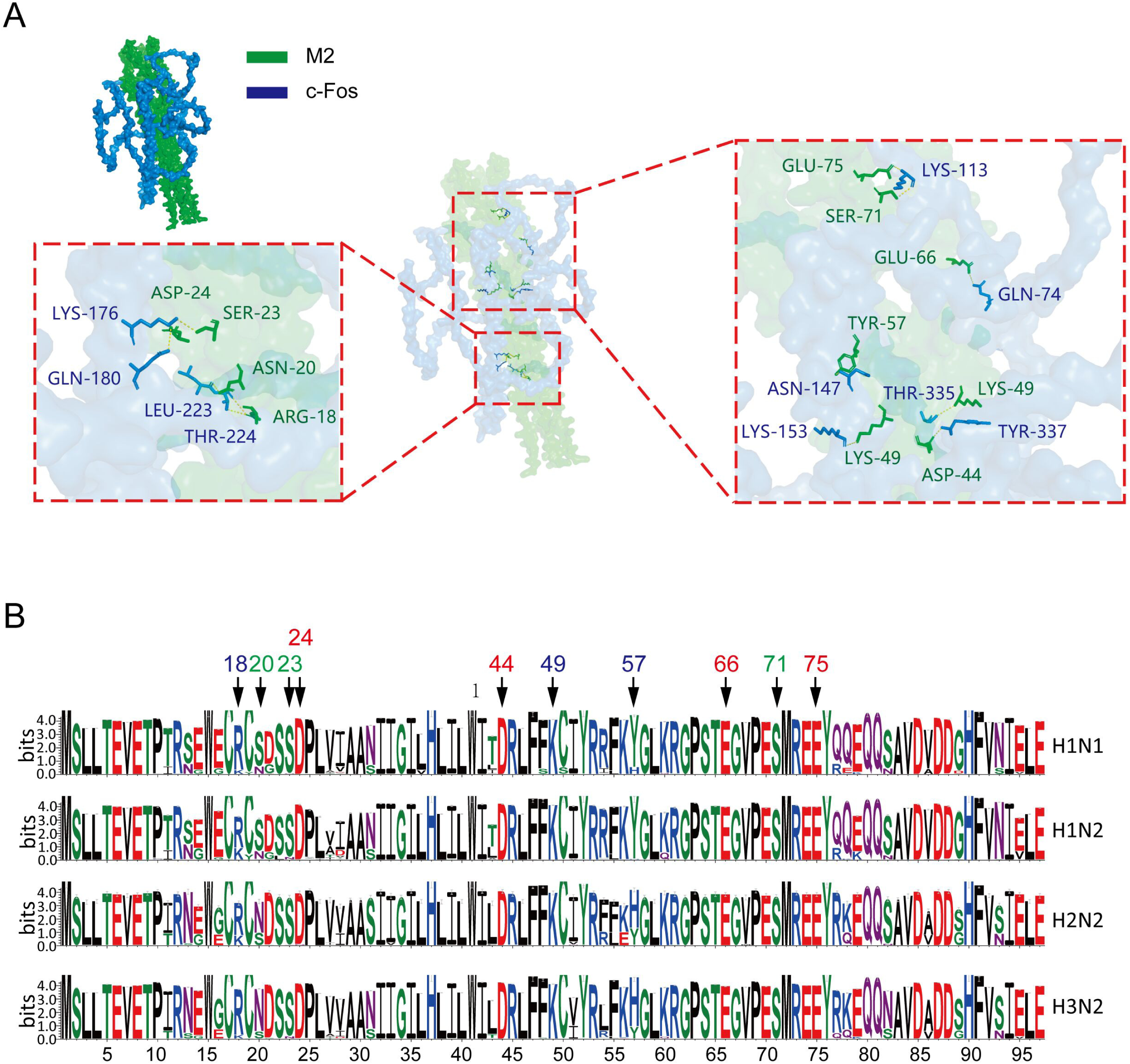
Predicted interaction sites of c-Fos-M2 and alignment of M2 sequences. (A) Molecular docking predicts potential interaction sites between c-Fos (in blue) and M2 (in green). c-Fos forms stable hydrogen bonds with the ARG-18, ASN-20, SER-23, ASP-24, ASP-44, LYS-49, TYR-57, GLU-66, SER-71, and GLU-75 sites of M2. (B) M2 protein amino-acid sequence logo of IAV isolates. The interaction sites of c-Fos-M2 are indicated by arrows. Logos were generated using the WebLogo3 online tool.

Given the functional importance and evolutionary conservation of these sites, designing drugs to disrupt c-Fos-M2 interaction could represent a promising strategy for developing effective anti-IAV therapeutics.

## DISCUSSION

IAV poses a significant threat to human health by triggering severe respiratory inflammation that can progress to acute lung injury and mortality. Understanding virus-host protein interactions is therefore critical for uncovering novel antiviral strategies. Our study elucidates a key mechanism centered on the host transcription factor c-Fos. We identify c-Fos as a proviral regulator that enhances IAV replication through a positive feedback loop by which IAV infection induces M2-mediated calcium influx, upregulating c-Fos, which in turn stabilizes M2 protein and promotes autophagosome accumulation, thereby amplifying viral propagation (Fig. 9). These findings not only delineate a novel pathway in IAV pathogenesis but also nominate c-Fos and its associated interactions as promising therapeutic targets.

**Fig 9.**
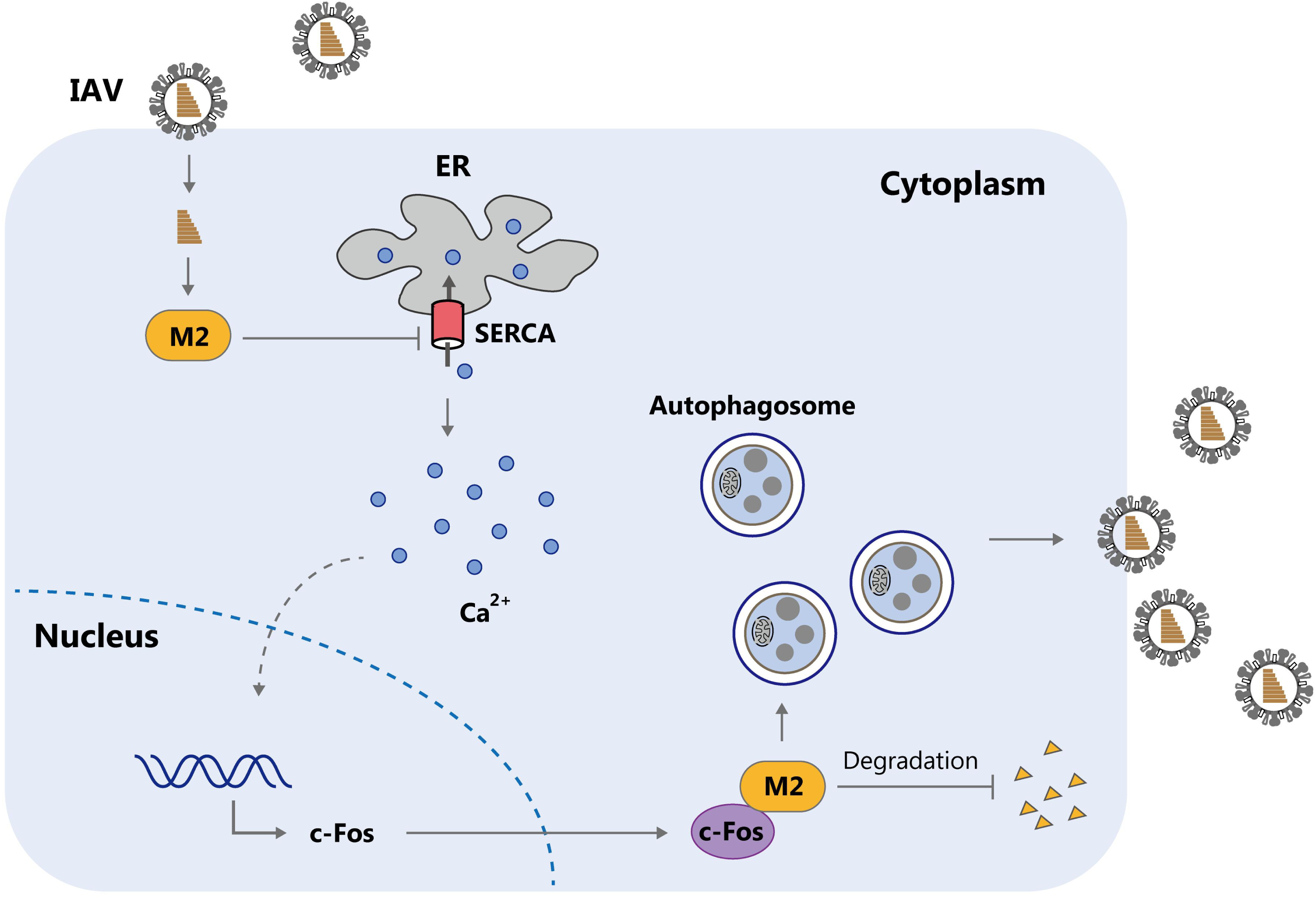
Schematic diagram of a model for the mechanism by which c-Fos promote IAV replication. Following influenza virus infection, the viral M2 protein inhibits the host SERCA pump, resulting in disrupted calcium homeostasis and elevated cytosolic calcium. This calcium elevation induces c-Fos upregulation, which consequently interacts with M2 and inhibits its degradation. The stabilized M2 thereby promotes virion assembly and budding, while also facilitating autophagosome accumulation to create a favorable intracellular environment for viral replication.

M2 protein serves critical roles in viral uncoating, assembly, and budding [5–7, 24, 25]. Therefore, the M2 protein facilitates IAV replication through multiple mechanisms. Stabilization of M2 protein in virus-producing cells enhances IAV replication efficiency. While c-Fos is traditionally known as a transcriptional regulator involved in fundamental cellular processes such as proliferation, differentiation, and apoptosis [10, 26, 27], we discovered it promotes IAV replication through a novel transcription-independent mechanism. Specifically, c-Fos physically interacts with the viral M2 protein, protecting it from both lysosomal and proteasomal degradation pathways. This interaction-mediated stabilization of M2 is crucial for optimal viral replication.

Our study further elucidates the complex interplay between IAV infection and autophagy. While it is well-established that IAV infection both induces autophagy initiation and inhibits autophagosome maturation [28], with the M2 protein being particularly implicated in autophagosome accumulation [8, 29–31], we identified the c-Fos-M2 axis as a key regulator of autophagosome accumulation that enhances viral replication. We demonstrated that M2 alone can induce LC3-II accumulation (Fig. 7A), an effect that is significantly amplified by c-Fos-mediated M2 stabilization (Fig. 7B and 7D). Notably, we found that c-Fos regulates autophagosome accumulation through additional mechanisms beyond M2 stabilization. Knockdown of c-Fos enhanced autophagic flux and promoted autophagosome-lysosome fusion even in uninfected cells (Fig. 3C), revealing its multifaceted role in autophagy regulation. While previous studies have shown c-Fos can induce autophagy through upregulation of BECN1 [32, 33], supporting our findings, the precise mechanisms governing c-Fos-mediated autophagy regulation in IAV-susceptible pulmonary cells require further investigation due to differences in experimental models.

Ca^2+^ is a ubiquitous second messenger that regulates diverse physiological processes, including cell survival, proliferation, apoptosis, and autophagy, through interactions with downstream effector proteins [34, 35]. Notably, Ca^2+^ signaling also plays critical roles in viral infections, particularly in IAV replication [36, 37]. Previous studies have shown that Ca^2+^-calmodulin complexes contribute to transcriptional regulation during IAV infection [38], while inhibition of Ca^2+^ channels disrupts progeny virion assembly [39]. While c-Fos expression is typically regulated by multiple signaling pathways and transcription factors, its regulation during IAV infection remained unclear. Building upon our finding that SERCA regulates c-Fos expression (with SERCA inhibition enhancing IAV-induced c-Fos upregulation), and given SERCA’s role as the ER calcium pump that maintains cytosolic Ca^2+^ homeostasis, and previous reports demonstrating that elevated cytosolic Ca^2+^ promote c-Fos transcription [32, 40]. We hypothesized that IAV modulates cytosolic calcium levels to drive c-Fos expression. Our study provides compelling evidence that IAV upregulates c-Fos expression by increasing cytosolic Ca^2+^ levels. Specifically, we demonstrated that M2-mediated elevation of cytosolic Ca^2+^ is pivotal for c-Fos upregulation (Fig. 4), and the Ca^2+^ chelator BAPTA-AM suppresses c-Fos expression, autophagy, and viral replication (Figs. 4-5). These findings align with established Ca^2+^-dependent c-Fos induction mechanisms while extending this paradigm to viral infection. Importantly, we reveal that cytosolic Ca^2+^ promotes IAV replication through c-Fos upregulation, demonstrating how calcium signaling modulates viral replication.

While the degradation pathways of IAV M2 protein have been partially characterized, current understanding remains incomplete. Previous studies have established that M2 undergoes lysosomal degradation in HeLa cells, mediated by the host factor MARCH8 [41]. However, our investigation reveals a more complex degradation mechanism that M2 is subject to dual regulation through both proteasomal and lysosomal pathways (Fig 6G and 6H). This expanded understanding likely reflects cell-type specific regulation, as our experiments were conducted in 293H cell line, distinct from the HeLa cell model used in prior studies. Most significantly, we identify the cellular transcription factor c-Fos as a novel regulator of M2 stability. Although c-Fos is canonically recognized for its transcriptional functions, our data uncover an important transcription-independent role in viral replication. Through comprehensive analyses, we show that c-Fos interacts with M2 protein and remarkably inhibits both proteasomal and lysosomal degradation pathways (Fig 6I). This dual inhibitory mechanism substantially prolongs M2 half-life, representing a sophisticated viral strategy to maintain critical M2 protein levels throughout infection. Our findings provide important insights into the host-virus interplay governing M2 protein stability. Given that M2 is essential for viral uncoating and assembly, its stabilization by c-Fos may represent a key strategy employed by IAV to optimize replication efficiency. Furthermore, the identification of c-Fos as a regulator of M2 degradation opens new avenues for antiviral research, as targeting the c-Fos-M2 interaction could potentially disrupt viral propagation.

Current influenza antivirals primarily target viral proteins, including neuraminidase inhibitors (e.g., oseltamivir, zanamivir), polymerase inhibitors (e.g., baloxavir marboxil), and M2 ion channel blockers (e.g., amantadine, rimantadine) [42–45]. However, the emergence of varying degrees of drug resistance to these agents [46–49] underscores the urgent need to identify novel antiviral targets through deeper investigation of IAV-host interactions. Our study demonstrates that IAV co-opts the host transcription factor c-Fos to stabilize viral M2 protein, thereby facilitating viral replication. This discovery reveals that disrupting the c-Fos-M2 interaction represents a promising antiviral strategy. Importantly, we predicted ten conserved M2 residues (ARG-18, ASN-20, SER-23, ASP-24, ASP-44, LYS-49, TYR-57, GLU-66, SER-71, and GLU-75) critical for this interaction, among which SER-23, ASP-24, ASP-44, LYS-49, GLU-66, SER-71, and GLU-75 exhibited pronounced evolutionary conservation across major human-infecting IAV subtypes (H1N1, H1N2, H2N2, and H3N2) (Fig. 8).The essential functional role and high conservation of these residues make the c-Fos-M2 interface an attractive target for developing broad-spectrum anti-influenza therapeutics with a high barrier to resistance.

In conclusion, our work establishes c-Fos as a critical host factor that supports IAV replication through M2 stabilization and autophagy promotion. Small-molecule inhibitors disrupting c-Fos-M2 binding or modulators of calcium/c-Fos signaling could represent promising candidates for future antiviral development.

## MATERIALS AND METHODS

### Cell lines, viruses and Mice strain

The human lung cancer cell lines NCI-H1395 (abbreviated as H1395) were cultured in RPMI 1640 medium (C11875500BT; Gibco) containing 10% fetal bovine serum (10270-106; Gibco) at 37°C and 5% CO_2_; 293H cells (used in previous studies and preserved in our laboratory) were cultured in Dulbecco’s modified Eagle medium (DMEM; C11995500BT; Gibco) containing 10% fetal bovine serum at 37°C and 5% CO_2_. The viral strains PR8 (A/Puerto Rico/8/1934) and WSN (A/WSN/1933) were preserved in our laboratory. Mice strain used in this study was BALB/c.

### Antibodies, reagents, plasmids, and short interfering RNA (siRNA) oligonucleotides

The following primary antibodies were used: anti-LC3B (2775; Cell Signaling Technology, CST), anti-SQSTM1 (5114; CST), anti-GAPDH (2118; CST), anti-NP (10780-01; SouthernBiotech), anti-SERCA (A1097; ABclonal), anti-Flag (AE092; ABclonal), and anti-influenza A virus M2 (IT-003-015; Immune Technology). Horseradish peroxidase (HRP)-conjugated goat anti-rabbit (E030120-01; EarthOx) and anti-mouse (E030110-01; EarthOx) secondary antibodies were also employed. Key reagents included CDN1163 (HY-101455; MedChemExpress), BAPTA-AM (HY-100545; MedChemExpress), T-5224 (B4664; APExBIO), cycloheximide (CHX, HY-12320; MedChemExpress), rapamycin (9904S; CST), MG132 (S2619; Selleck), bafilomycin A1 (A8510; Solarbio), and chloroquine (C6628; Sigma). siRNA oligonucleotides targeting ATP2A2 (5′-GGTGCTATTTACTACTTTA-3′), c-Fos (5’-TCTGCTTTGCAGACCGAGATT-3’; 5’-GTGGAACAGTTAUCTCCAGAA-3’ and 5’-CACTGCTTACACGTCTTCCTT-3’), and a negative control siNC (5′-TTCTCCGAACGTGTCACGT-3′) were synthesized by Synbio Technologies. Plasmids for mammalian expression of influenza A virus genes (PB2, PB1, PA, HA, NP, NA, M1, M2, NS1, NS2) were preserved in our laboratory.

### RNA-Seq Analysis

Following total RNA extraction, mRNA was enriched using oligo(dT)-conjugated magnetic beads and subsequently fragmented at elevated temperature. The fragmented mRNA served as a template for first-strand cDNA synthesis via reverse transcription. During second-strand synthesis, the cDNA underwent end repair and adenine (A)-tailing. Sequencing adapters were then ligated to the fragments, which were purified and size-selected using selection beads. Finally, the library was amplified via PCR and sequenced on an Illumina NovaSeq 6000 platform.

### Real-time quantitative PCR (RT-qPCR) analysis

Total RNA from H1395 cells was extracted with the Universal RNA Purification Kit (EZB-RN4; EZBioscience) and reverse-transcribed into cDNA using HiScript III RT SuperMix. qPCR was performed using a 2 × SYBR Green Master Mix (A0001; EZBioscience) on a LightCycler96 system (Roche) with the following protocol: 95°C for 30 sec, then 40 cycles of 95°C for 10 sec and 60°C for 30 sec. Relative mRNA expression was normalized to GAPDH and calculated using the 2^–ΔΔCT method.

### Plaque assay

Viral titers were determined by plaque assay. Briefly, confluent MDCK monolayers in 12-well plates were inoculated with serially diluted viral supernatants (0.25 mL) and incubated at room temperature for 1 h with gentle swirling every 15 min. After the incubation, the cells were overlaid with 1 mL of culture medium containing 1% agarose and allowed to solidify at room temperature. The plates were then inverted and incubated at 37°C for 72 h. Finally, the cells were fixed and stained with 0.1% crystal violet after removal of the agarose overlay to visualize the plaques.

### Immunofluorescence assay and Confocal Fluorescence Microscopy

Following treatments, cells were fixed, permeabilized, and blocked. They were then incubated with primary antibodies at 4°C overnight, followed by species-specific secondary antibodies (Alexa Fluor 488 or 594) and DAPI for nuclear staining. Images were acquired using an Olympus FV10i confocal microscope with sequential scanning. The number of LC3B puncta per cell was quantified blindly using ImageJ software.

### Flow Cytometry

To measure cytosolic Ca^2+^ levels in H1395 and 293H cells, we used the fluorescent Ca^2+^ indicator Fluo-4 AM (F312, Dojindo). Cells were washed three times with PBS and loaded with 5 μM Fluo-4 AM for 30 minutes. After incubation, the cells were harvested by centrifugation and analyzed via flow cytometry. For each sample, 30,000 events were recorded and used for subsequent analysis.

### Co-immunoprecipitation (Co-IP)

Cells in culture were washed twice with ice-cold PBS to completely remove residual medium. IP lysis buffer supplemented with protease inhibitor (Beyotime, ST507) was directly added onto cell layers and scraped on ice. Cell lysates were processed as mentioned above. For immunoprecipitation of FLAG-tagged proteins, cell lysates were incubated with anti-FLAG magnetic beads (MedChemExpress, HY-K0207) on a rotator at 4°C overnight. IP beads were then washed with IP lysis buffer four times, and boiled in 1 × SDS loading buffer at 98°C for 5 min for SDS-PAGE and immunoblot analysis.

### Western blotting

Cells were lysed on ice and centrifuged to collect supernatants. Protein concentrations were determined using a BCA assay kit (23225; Thermo Scientific). After normalization, samples were boiled in SDS loading buffer, separated by SDS-PAGE, and transferred to PVDF membranes (IPVH00010; Millipore). The membranes were blocked with 5% non-fat milk and incubated with primary antibodies overnight at 4°C, followed by HRP-conjugated secondary antibodies at 37°C for 2 h. Protein bands were visualized using Clarity Western ECL substrate (170-5060; Bio-Rad).

### Cell viability assay

Cell viability was evaluated using cell counting kit-8 (CK04; Dojindo) 24 h after H1395 cell monolayers were exposed to different concentrations of BAPTA-AM and T5224 according to the manufacturer’s instructions and compared with negative controls. The optical density at 450 nm was used as a measure indicating cell viability and was determined using a microplate spectrophotometer (BioTek).

### Protein degradation assay

293H cell lines were cotransfected with M2 plasmid with empty vector or Flag-c-Fos plasmid. After 24 h post-transfection, CHX (50 μg/mL) was added to the medium and cells were harvested at different time points. Western blotting of cell lysates was performed to detect Flag-c-Fos, M2 and GAPDH. To investigate the cellular pathway of M2 degradation, MG132 (200 nM) or Bafimycin A1 (2 nM) was added to the medium, and the cells were harvested 24 h after treatment. Three independent experiments were performed.

### Molecular docking analysis

The structure of of c-Fos and M2 proteins were retrieved from the AlphaFold Protein Structure Database [50]. Using Gramm-X [51] (https://gramm.compbio.ku.edu/), we performed rigid docking analysis of c-Fos and M2 to obtain the top 10 docking models. The highest-ranked model was visualized using PyMOL software.

### Selection of M2 sequences for alignment

The M2 protein sequences of human IAVs were retrieved from the UniProt database. Strain selection was refined by removing duplicate M2 sequences and retaining only those with complete coverage. Additionally, the HA/NA subtypes of these strains were identified. Based on these criteria, a curated list of the most representative M2 sequences was compiled. These sequences were then aligned using MAFFT, and the resulting multiple sequence alignment was analyzed with the WebLogo3 online tool to generate a sequence logo.

### Statistical analysis

Data in this study are expressed as the means ± standard deviations (SD), and differences were measured using a Student’s t test. A P value of <0.05 was considered statistically significant. For all analyses, the notations used to indicate significance between groups are the following: *, P < 0.05; **, P <0.01; ***, P <0.001; #, P >0.05.

## ETHICS STATEMENT

All animal experiments were approved by the Animal Ethics Committee of Guangzhou Medical University (Approval No. 20240954) and were conducted in compliance with the NIH Guide for the Care and Use of Laboratory Animals.

## ACKNOWLEDGEMENTS

This work was supported by the Basic and Applied Basic Research Foundation of Guangdong (Grant No. 2025A1515012645); the Open Research Project of the Key Laboratory of Viral Pathogenesis & Infection Prevention and Control of the Ministry of Education (Grant No. 2023VPPC-R04); and the Basic and Applied Basic Research Foundation of Guangdong (Grant No. 2025A1515012487).

## SUPPLEMENTAL MATERIAL

S1 Table. Primers used for RT-qPCR.

## DATA AVAILABILITY

All relevant data are within the Manuscript, Figures and Supplemental material files. The RNA-Seq data have been deposited in GSA public database under the accession number HRA014120.

